# Structural basis for impaired oxygen evolution in extrinsic-protein-reconstituted photosystem II

**DOI:** 10.64898/2026.01.11.698909

**Authors:** Yoshiki Nakajima, Koji Kato, Jian-Ren Shen, Ryo Nagao

## Abstract

Photosystem II (PSII) is a membrane-bound pigment-protein complex in oxygenic photosynthesis that catalyzes water splitting and oxygen evolution. Here, we present the X-ray crystallographic structure of a PsbO/V/U-reconstituted PSII from *Thermosynechococcus vulcanus* at 2.0 Å resolution, revealing proper rebinding of the three extrinsic subunits, PsbO, PsbV, and PsbU. The overall geometry of the Mn_4_CaO_5_ cluster is largely preserved, although a subtle shortening of the Mn2–O2 bond suggests a minor local rearrangement. Structural analysis identifies perturbations that may underlie the reduced oxygen-evolving activity, including altered bicarbonate-binding interactions on the electron acceptor side and the loss of water molecules W658 and W660 in the O1 channel, disrupting a hydrogen-bond network critical for water delivery. In contrast, the Cl-1 and O4 channels remain intact. These findings suggest that disrupted water delivery and electron transport, together with minor rearrangements within the Mn_4_CaO_5_ cluster, may contribute to the decreased activity of the reconstituted PSII.

## Introduction

Oxygenic photosynthetic reaction by cyanobacteria, algae, and land plants converts solar energy into biologically useful chemical energy concomitant with evolution of molecular oxygen (*1*). The photosynthetic oxygen evolution is performed by a dimeric, multi-subunit membrane protein complex of photosystem II (PSII) (*2–6*), whose monomeric unit in cyanobacteria is composed of 17 transmembrane subunits and three extrinsic subunits in addition to more than 80 cofactors, which include 35 chlorophylls *a* (Chls), two pheophytins *a* (Pheos), 12 carotenoids (Cars), two plastoquinones (PQs), one Mn_4_CaO_5_ cluster, two hemes, one non-heme iron, one bicarbonate, two chlorides, and 25 lipids, as revealed by its X-ray crystal structure (*4, 7, 8*).

The central part of PSII is the D1 and D2 subunits, which possess redox cofactors responsible for electron-transfer reactions (*2–8*). Following light harvesting and excitation-energy transfer in PSII, a charge separation occurs between P680 and Pheo_D1_, forming the P680^+^Pheo_D1_^−^ species (*9, 10*). On the electron acceptor side (stromal/cytosolic side), Pheo_D1_ acts as an electron acceptor, whereas Pheo_D2_ is redox-inactive (*11*). The electron on Pheo_D1_^−^ is transferred to the primary quinone electron acceptor Q_A_ and then to the secondary quinone electron acceptor Q_B_ (*12–14*). On the electron donor side (lumenal side), P680 is composed of dimeric Chls, P_D1_ and P_D2_, and forms the rection center Chls together with two monomeric Chls, Chl_D1_ and Chl_D2_. The initial charge separation occurs mainly on Chl_D1_ (*9, 15, 16*), and finally a positive charge is located mostly on P_D1_ (*17–20*). The positive charge on P680^+^ is reduced by the Mn_4_CaO_5_ cluster through a redox-active tyrosine Y_Z_ (D1-Y161) located between the Mn_4_CaO_5_ cluster and P680 (*7, 8, 21*). The atoms in the Mn_4_CaO_5_ cluster are designated as Ca, Mn1–Mn4, and O1–O5 (*7*). Another redox-active tyrosine Y_D_ (D2-Y160) functions as a peripheral electron donor to P680^+^ (*22*).

The oxygen-evolving reaction takes place in the Mn_4_CaO_5_ cluster at the oxygen-evolving complex (OEC) (*3–8*), which proceeds through a light-driven cycle of five intermediates called S states (S_0_**–**S_4_) (*2–6, 23*). The S_1_-state is the most stable one in the dark (*23*), which proceeds to the next S*_n+1_* state (*n* = 0**–**3) upon one-electron oxidation, except that the most oxidized S_4_-state is unstable and immediately relaxes to the S_0_-state by releasing an O_2_ molecule (*3–6*). During this S-state cycle, two water molecules are split, concomitant with the release of four protons. The Mn_4_CaO_5_ cluster is stabilized by amino-acid ligands and water molecules (*4, 7, 8*), and the structural environment of the Mn_4_CaO_5_ cluster is also maintained by extrinsic subunits of PSII. The PSII extrinsic subunits are largely diversified from cyanobacteria to land plants, although the membrane-spanning PSII-core subunits are mostly conserved among oxyphototrophs (*24, 25*). The molecular diversity and evolution of the extrinsic subunits of PSII have been reported extensively (*26–30*).

Mutagenesis and biochemical studies have been performed to study the effects of extrinsic proteins on the oxygen-evolving reaction of PSII (*26–30*). In particular, release-reconstitution experiments of extrinsic proteins have been used for a long time to reveal their functional and binding properties, which were mainly assessed by oxygen-evolving activity of the PSII samples reconstituted with extrinsic subunits (*26–28*). However, the activity does not recover fully after reconstitution of all the extrinsic proteins to the PSII samples from any oxyphototrophs (*26–28*). This may be because (i) the extrinsic proteins cannot bind to their proper sites at the lumenal side of PSII by the release-reconstitution experiments and/or (ii) structural changes in the OEC occur by the release-reconstitution experiments albeit with their proper bindings.

The cyanobacterium *Thermosynechococcus vulcanus* NIES-2134 is used for biochemical and structural studies of PSII (*4*). The characteristics of the *T. vulcanus* PSII are highly active and stable, thereby enabling us to make high-quality crystals for X-ray crystallography (*31, 32*). The *T. vulcanus* PSII contains three types of extrinsic subunits, PsbO, PsbV, and PsbU (*7, 33*). The oxygen-evolving activity of the PsbO/V/U-reconstituted PSII showed 80–87% recovery compared with native PSII cores (*33, 34*), and it is not clear why 100% recovery of the activity is not achieved. The X-ray crystallographic analysis of the PsbO/V/U-reconstituted PSII from *T. vulcanus* may address these issues as described above (labeled (i) and (ii)).

In this study, we determined an X-ray crystal structure of the PsbO/V/U-reconstituted PSII prepared from *T. vulcanus*. The result clearly revealed that all the three extrinsic subunits, PsbO, PsbV, and PsbU, are properly rebound to their native positions on the PSII core. In the PsbO/V/U-reconstituted PSII, however, it is found that not only the bicarbonate structure is altered but also some water molecules in the OEC are lost, which may be the reason why the activity is not recovered to 100%.

## Results and discussion

### Overall structure of the PsbO/V/U-reconstituted PSII

The release and reconstitution of the three extrinsic proteins were confirmed by SDS-PAGE (Fig. S1), which showed their complete removal from the PSII cores (lane 2) and successful reconstitution (lane 3) relative to the untreated PSII (lane 1). The 13–20% decrease in O₂ evolution following reconstitution has been consistently reported previously (*33, 34*), and is referenced here as an established characteristic of the system.

The crystal structure of PsbO/V/U-reconstituted PSII was solved at a resolution of 2.0 Å (Table S1). The structure clearly shows associations of the three extrinsic subunits with PSII cores (Fig. 1A), and is well fitted into the native PSII structure (PDB: 3WU2) (Fig. 1B). As observed in previous PSII crystal structures (*35, 36*), only one monomer of the dimeric PSII retained the PsbY subunit in the present structure (Fig. 1B). In this study, we define the monomer containing PsbY as the A-monomer, and the other as the B-monomer (Fig. 1B). Unless otherwise stated, all structural comparisons are made between the A-monomers. Detailed comparisons of individual extrinsic proteins revealed that only the interaction of PsbO-R60 with CP47-G404 and that of PsbV-K124 with PsbE-R61 differ between the reconstituted and native PSII structures, where the hydrogen bonds at distances of 3.1 Å formed between these amino acid pairs in the native PSII are broken in the reconstituted PSII (Fig. 1C–G). In addition, the PsbO/V/U-reconstituted PSII retains all Chl and Car molecules in both monomers (Fig. S2A, B) and depicts no substantial changes in the distances among the redox-active cofactors including the Mn_4_CaO_5_ cluster (Fig. S2C). The root mean square deviation (RMSD) of the structures between reconstituted and native PSII (PDB code: 3WU2) is 0.172 Å for 5243 Cα atoms. These results indicate that no dramatic structural changes occurred in the PsbO/V/U-reconstituted PSII.

**Figure 1.**
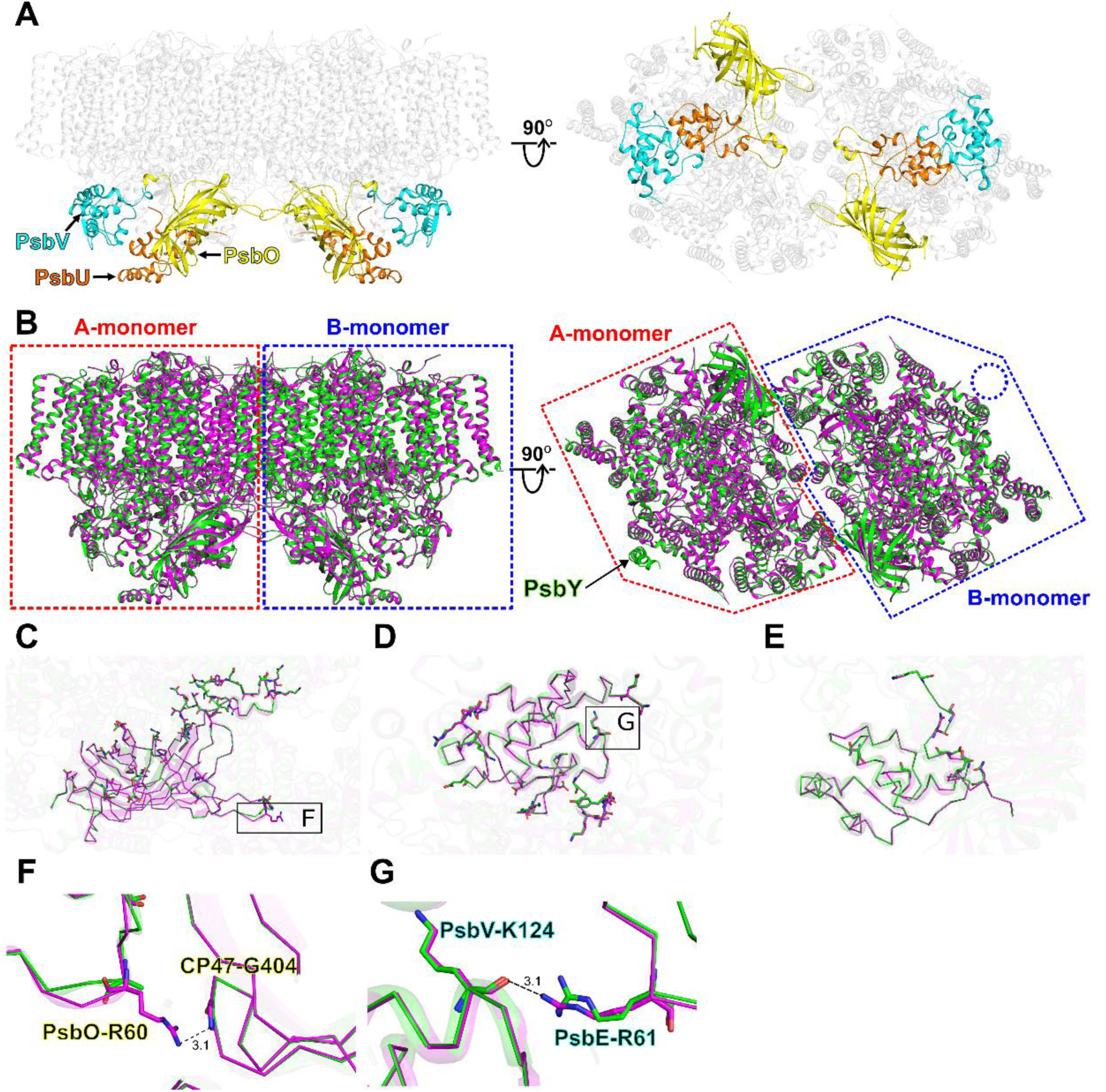
Structure of the PsbO/V/U-reconstituted PSII. (**A**) Structures of the reconstituted PSII are viewed from the direction perpendicular to the membrane normal (left panels) and the lumenal side (right panels). Only protein structures are shown, and cofactors are omitted for clarity. The extrinsic proteins are colored yellow (PsbO), cyan (PsbV), and orange (PsbU), whereas the intrinsic proteins are colored grey. (**B**) Superposition of the PsbO/V/U-reconstituted PSII (green) with the native PSII structure (magenta; PDB: 3WU2). The red and blue dashed lines indicate the A- and B-monomers, respectively, as defined in this study. (**C–G**) Detailed structural comparisons of PsbO (**C, F**), PsbV (**D, G**), and PsbU (**E**) between the reconstituted and native PSII structures. The areas encircled by black squares in panels **C** and **D** are enlarged in panels **F** and **G**, respectively. Interactions are indicated by dashed lines, and the numbers are distances in Å.

To further assess whether the release and reconstitution of the extrinsic proteins affected the OEC, we evaluated the interatomic distances, in both the Mn_4_CaO_5_ cluster, as well as those between the cluster and its amino acid ligands between the native and reconstituted PSII structures. The average deviation of interatomic distances between the two structures was calculated to be 0.132 Å (Table S2). The largest difference was observed for the Mn1–O1 pair, which exhibited an elongation of 0.13 Å in the reconstituted PSII (reliability = 67.2%; calculated using 0.132 Å as one standard deviation; see Methods; Table S3). This variation, however, is not statistically significant, indicating that the detachment and reassociation of the extrinsic subunits exert only a minimal effect on the OEC geometry. Consistently, the overall architecture of the Mn cluster remained virtually unchanged between the two structures (Fig. 2, Fig. S3). At a finer level of analysis, however, a closer inspection of interatomic distances revealed subtle but quantifiable local differences within the OEC, which could influence the microenvironment of the catalytic site.

**Figure 2.**
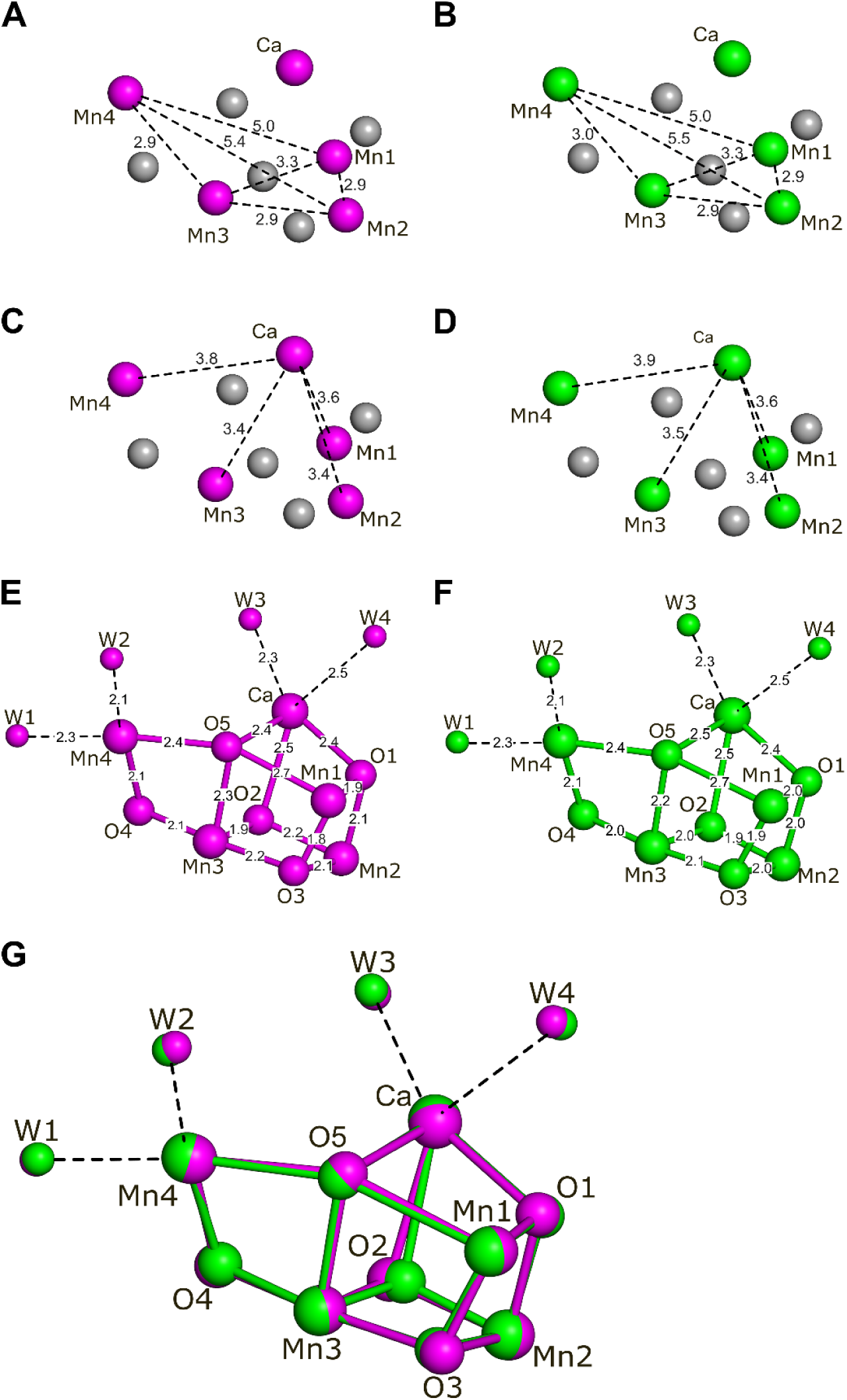
Structural comparison of the OEC in the A-monomers of the native and reconstituted PSII. (**A, B**) Distances between Mn atoms. (**C, D**) Distances between Mn and Ca atoms. (**E, F**) Distances between the Mn_4_CaO_5_ cluster and its coordinating water molecules. (**G**) Superposition of the OECs from the two PSII structures. All panels show the OECs in the A-monomers of the native PSII (magenta; PDB 3WU2) and the PsbO/V/U-reconstituted PSII (green).

### Structural comparison of the OEC

To characterize these local variations in detail, we analyzed interatomic and ligand distances between the native and reconstituted PSII structures. The average deviation of interatomic distances between the two structures was 0.132 Å (Table S2). Among all pairs, the Mn2–O2 bond showed the largest difference, displaying a 0.26 Å shortening in the reconstituted PSII relative to the native structure (reliability = 95.1%, 1.97σ assuming σ = 0.132 Å; see Methods; Tables S2, S3). This deviation exceeds the coordinate error and therefore represents a statistically significant change.

Structural superposition of the Mn_4_CaO_5_ clusters confirmed a subtle displacement corresponding to this bond shortening (Fig. 2). However, the Mn2–O2 bond is known to exhibit substantial flexibility, as its length also varies considerably even between the A- and B-monomers of the native PSII. In line with this inherent mobility, comparison of the B-monomers between the two structures revealed no notable differences (Fig. S3). Taken together, these findings indicate that the observed Mn2–O2 shortening represents a statistically significant, localized structural rearrangement within the OEC. While this change does not imply a global alteration of the Mn_4_CaO_5_ cluster, it is likely to modulate the local coordination environment and may influence the stability and/or reactivity of the catalytic site.

### Structural environments around the bicarbonate

Despite the overall similarity of the structure between the reconstituted and native PSII, some structural changes are found in the bicarbonate molecule and its binding environment (Fig. 3A). The angle of the bicarbonate shifted 8.0 degrees toward the D1-Y246 side in the reconstituted PSII (Fig. 3B), when the structures are superimposed based on the non-heme iron between the reconstituted and native PSII. This angular shift was even more pronounced in the B-monomer, reaching 9.9 degrees (Fig. S4A, B).

**Figure 3.**
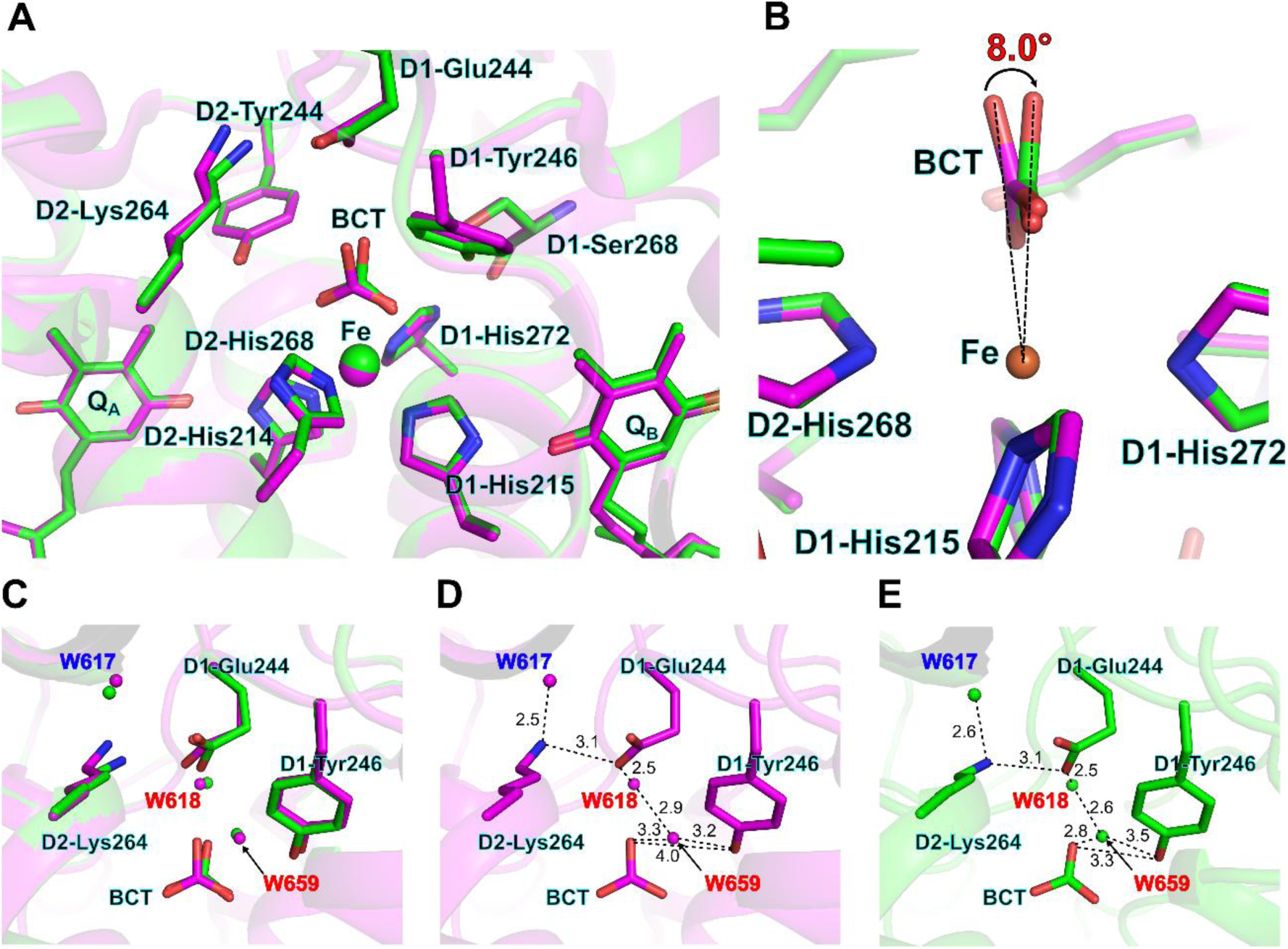
Structural comparison of the acceptor side in the A-monomers of the native and reconstituted PSII. (**A**) Structures around the bicarbonate (BCT) molecule and non-heme iron (Fe). The PsbO/V/U-reconstituted and native PSII structures are colored green and magenta, respectively. (**B**) Angle of BCT. The angle was calculated from three points: O1 of BCT in the native PSII (3WU2), Fe, and O1 of BCT in the reconstituted PSII, based on superposition of the Fe atom between the two structures. (**C–E**) Superposition of BCT and its nearby amino acid residues and water molecules between the structures of reconstituted and native PSII (**C**), structure of the reconstituted PSII (**D**), and structure of the native PSII (**E**). Interactions are indicated by dashed lines, and the numbers are distances in Å. Water molecules of the D1 and D2 subunits are labeled with red and blue colors, respectively.

To clarify whether the observed shift reflects a genuine rearrangement of the bicarbonate or merely a local distortion of the surrounding protein environment, we evaluated the distance between the O1 atom of the bicarbonate and the hydroxyl group of D1-S268 (Fig. S5). Although D1-S268 lies in close proximity to the bicarbonate, it does not form a direct hydrogen bond (Fig. S5) and therefore serves as a stable internal reference point. Consistent with this assumption, the distance between the non-heme iron and D1-S268 in the two PSII structures differed by 0.06 Å between the A-monomers (Fig. S5A, B) and 0.01 Å between the B-monomers (Fig. S5C, D), corresponding to 0.377 σ and 0.063 σ, respectively, based on the diffraction-component precision index (DPI)-derived error criterion (1 σ = 0.159 Å), indicating that D1-S268 itself remains structurally unchanged in the reconstituted PSII. In contrast, relative to this invariant residue, the distance between the bicarbonate O1 atom and D1-S268 increased by 0.58 Å in the A-monomers (Fig. S5A, B) and 0.61 Å in the B-monomers (Fig. S5C, D), corresponding to 3.65 σ and 3.84 σ, respectively, under the same criterion. These highly significant deviations indicate that the structural change observed in the reconstituted PSII arises from a genuine displacement of the bicarbonate molecule rather than from alterations in the surrounding protein environment.

The protein structure around the bicarbonate is also changed in the reconstituted PSII (Fig. 3C–E), where side chains of D1-E244, D1-Y246, and D2-K264 are shifted slightly toward the bicarbonate, although their backbones are virtually identical to the native PSII structure. In conjunction with the protein movement, the positions of water molecules, specifically W618 and W659 of D1, and W617 of D2, shifted their positions slightly relative to those in native PSII. As a result, the bicarbonate molecule is shifted toward W659 and D1-Y246 in the reconstituted PSII, with the distances between bicarbonate and W659/D1-Y246 shortening from 3.3/4.0 Å to 2.8/3.3 Å, respectively (Fig. 3D, E).

In the native PSII, no water molecule corresponding to W659 of D1 in the A-monomer was detected in the B-monomer (Fig. S4C–E). In this structure, the hydrogen-bond distances between the bicarbonate and W659/D1-Y246 are relatively long, suggesting weaker hydrogen bonds and less stable water binding (Fig. 3D). In contrast, in the reconstituted PSII, the bicarbonate exhibits altered coordination geometry accompanied by shortened hydrogen-bond distances, implying stronger hydrogen-bond interactions and the retention of water molecules at equivalent positions in both monomers. The presence of this water molecule in the B-monomer therefore provides structural evidence for a modified and more stable binding state of the bicarbonate in the reconstituted PSII.

D1-Y246 plays important roles in photoautotrophic growth and oxygen-evolving reactions (*37–40*), especially in electron transfer from Q_A_ to Q_B_ (*38, 40*), by stabilizing the bicarbonate molecule as well as water environments (*13, 39–41*). The mutation of D1-Y246 gave rise to both an enhancement of oxygen-evolving activity with supplementation of bicarbonate and a decrease in the redox potential of Q_B_ (*40*). Additionally, theoretical calculations suggested both a different pathway participated by water molecules and destabilization of the bicarbonate molecule upon the D1-Y246 mutation (*40*). Here, we clearly show alterations in the hydrogen-bond interactions formed between D1-Y246 and bicarbonate in the reconstituted PSII (Fig. 3C–E). Since the PsbO/V/U-reconstituted PSII showed a decrease in oxygen-evolving activity and alteration in the thermoluminescence glow curves compared with native PSII (*33, 34*), the structural changes around the bicarbonate observed in the reconstituted PSII structure are likely attributable to the retardation of electron transfer from Q_A_ to Q_B_.

The position of D1-E244 is shifted also in the PsbO/V/U-reconstituted PSII structure (Fig. 3C–E). It was suggested that replacement of D1-E244 with Ala induces structural alterations, resulting in a major impact on the Q_B_ binding site (*38*). Both structures of Q_A_ and Q_B_ remain almost the same between the native and reconstituted PSII, implying that changes in the D1-E244 residue may have induced reductions in the oxygen-evolving activity upon release and reconstitution of the extrinsic proteins.

Two tyrosine residues, D1-Y246 and D2-Y244, are symmetrically positioned at the bicarbonate-binding site within hydrogen-bonding distances (*7, 8, 13*). Like D1-Y246, D2-Y244 plays a role in supporting photoautotrophic growth and oxygen-evolving activity, suggesting a contribution of D2-Y244 to the stabilization of the bicarbonate binding and electron transfer from Q_A_ to Q_B_ (*40, 42*). However, D2-Y244 does not appear to shift significantly in the PsbO/V/U-reconstituted PSII structure (Fig. 3A). These observations suggest that D1-Y246, rather than D2-Y244, is more critical for stabilizing bicarbonate, influencing the Q_B_ structure and the surrounding water molecules, in the reconstituted PSII.

Removal and binding of extrinsic proteins have been reported to affect the acceptor side of PSII. Kato and Noguchi used FTIR spectroelectrochemistry to analyze changes in the Q_A_ redox potential in spinach PSII after salt-induced removal of the extrinsic proteins PsbO, PsbP, and PsbQ (*43*). This study revealed that removal of extrinsic proteins from the lumenal side can modulate the redox environment surrounding Q_A_ on the stromal side. Furthermore, Yamada et al. showed that binding of the red algal extrinsic protein of PsbQ′ to *T. elongatus* PSII altered the redox potential of Q_A_ (*44*), highlighting the impact of extrinsic-protein association on the acceptor side of PSII. These observations support the view that the structural and functional changes observed in this study reflect modulation of the acceptor side by the association or dissociation of extrinsic proteins.

### Loss of water molecules in the O1 channel

Three primary channels are implicated in putative proton and water transfer pathways around the Mn_4_CaO_5_ cluster: the Cl-1 channel, the O4 channel, and the O1 channel (*45–54*). Two water molecules observed in the O1 channel of native PSII, namely W660 of D1 and W658 of D1, disappeared in the PsbO/V/U-reconstituted PSII structure (Fig. 4A). In the native PSII structure, W660 interacts with D1-N298 at a distance of 2.5 Å and forms a hydrogen-bond network with D1-N298 as well as W562 of D1 and W697 of CP43 (Fig. 4B, C). However, in the reconstituted PSII structure, absence of W660 results in a shift of D1-N298, which now interacts with W562 and W697 at distances different from that of the native structure (Fig. 4B, D). This shift appears to preserve the hydrogen-bond interactions between D1-N298 and W562/W697.

**Figure 4.**
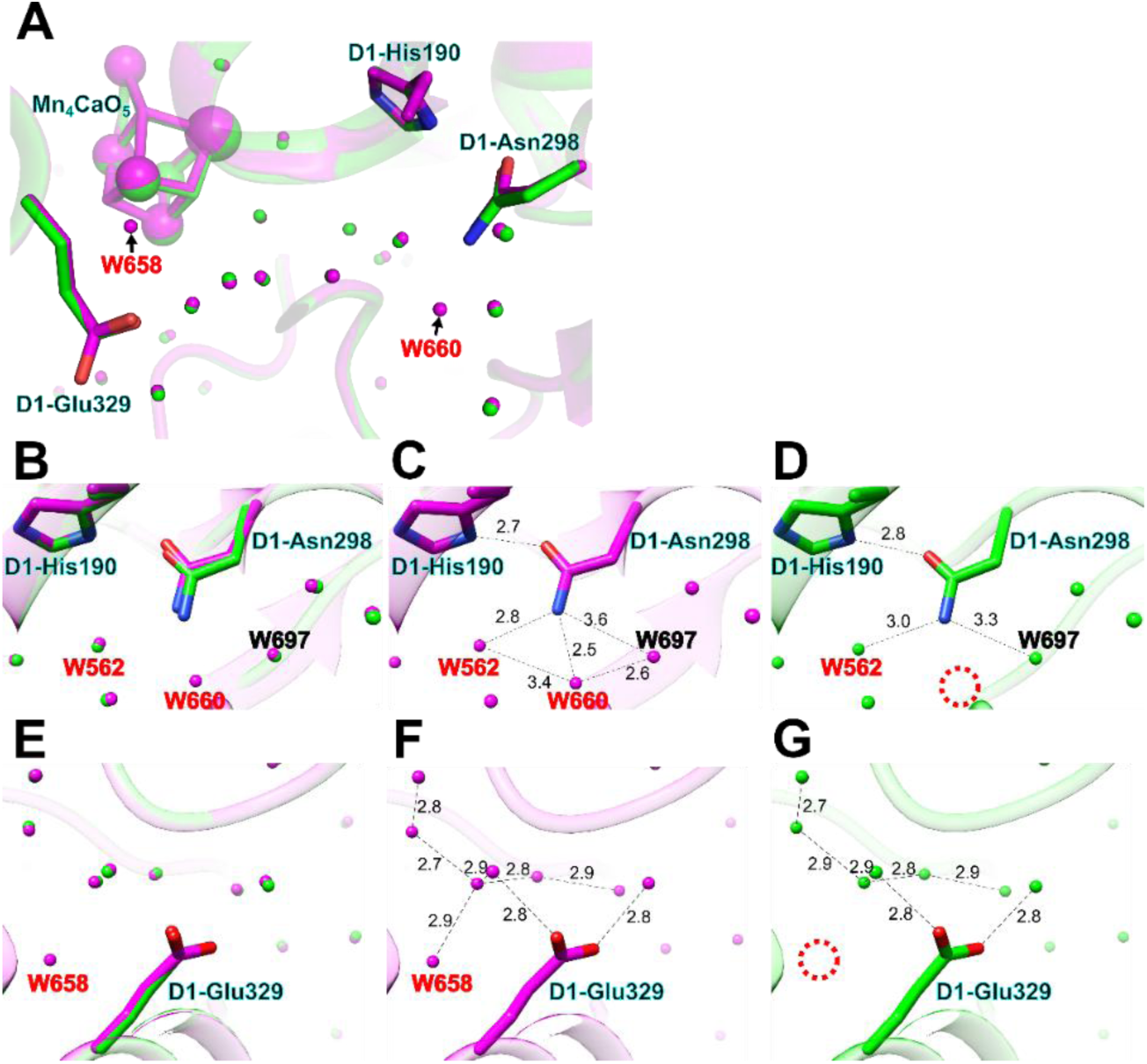
Structural comparison of the O1 channel in the A-monomers of the native and reconstituted PSII. (**A**) Structures of proteins and cofactors in the O1 channel. The PsbO/V/U-reconstituted and native PSII structures are colored green and magenta, respectively. (**B–D**) Interactions of W660 with proteins and water molecules in the structures of the reconstituted and native PSII superposed (**B**), the native PSII (**C**), and the reconstituted PSII (**D**). (**E–G**) Interactions of W658 with protein and water molecules in the structures of the reconstituted and native PSII superposed (**E**), the native PSII (**F**), and the reconstituted PSII (**G**). Interactions are indicated by dashed lines, and the numbers are distances in Å. Water molecules of the D1 and CP43 subunits are labeled with red and black colors, respectively. Areas enclosed by red dashed circles indicate the absence of W562 (**D**) and W658 (**G**) in the reconstituted PSII.

W658 forms a hydrogen-bond network across water molecules (Fig. 4E, F) and is positioned near D1-D342, D1-H332, and Mn1 in the native PSII structure (*7, 8*). However, W658 in the PsbO/V/U-reconstituted PSII disappears; instead, a cavity is formed at the site of W658 (Fig. 4E–G). Disappearance of W658 does not seem to affect the neighboring water molecules and amino acid residues, as there are no critical movements of such molecules in the reconstituted PSII in comparison with the native PSII (Fig. 4B–D).

The two water molecules are included in the O1 channel, which is a putative water delivery and/or proton transfer pathway from lumen to the Mn_4_CaO_5_ cluster, and is also called a large channel that starts from a site near Y_Z_ and passes through O1 and D1-N298 (*7, 45–51, 55*). This channel contains four minor contributing channels in total (*53*), one of which is the Y_Z_ network having a rigid hydrogen-bond network suitable for proton transfer (*50*). The serial femtosecond X-ray crystallography studies showed that water molecules in the O1 channel were significantly altered during the S_2_ → S_3_ transition (*56, 57*). The altered water molecules are located between O1 and D1-N298, and are designated as the "water wheel", which is likely to serve as an entry point for the substrate water (*57*). An Ala mutant of D1-N298 showed a decrease in the efficiency of the S_2_ → S_3_ transition and severely inhibited the S_3_ → S_0_ transition (*58*). Furthermore, time-resolved infrared (TRIR) analysis using this mutant with NO_3−_ substitution suggested that water delivery and proton release occur through the O1 and Cl-1 channels, respectively, during the S_2_ → S_3_ transition (*59*). Indeed, the Cl-1 channel has been suggested to function in proton release during the S_2_ → S_3_ and S_3_ → S_0_ transitions (*60–65*), and remarkable structural changes has been found during the S_2_ → S_3_ transition recently (*66*).

The absence of W660 in the PsbO/V/U-reconstituted PSII structure disrupts the hydrogen-bond network that are otherwise present in the native PSII structure. To compensate this network, the side chain of D1-N298 is moved toward W697 in the reconstituted PSII (Fig. 4B, D). This structural change may compromise the function of Y_Z_ due to the form of the D1-Y161(Y_Z_)-H190-N298 triad. In contrast, W658 is positioned near the Mn_4_CaO_5_ cluster and its surrounding amino acid ligands, implying its involvement in the structural stability of the water network around the Mn1 and O1 atoms of the Mn_4_CaO_5_ cluster, despite the fact that disappearance of W658 showed little effects on its surrounding water molecules at the dark-adapted S_1_ state. Based on the role of the O1 channel, it is suggested that both W660 and W658 participate in water delivery during the S_2_ → S_3_ transition; disappearance of these water molecules in the reconstituted PSII may result in the decrease of the oxygen-evolving activity.

The O1 channel is separated into two branches toward the lumenal side: one is from D1-E189/D342 to CP43-K79/N418 and PsbV-Q34 via CP43-L401/G409 as branch A, and the other one is from D1-E189/D342 to D2-E343 and PsbU-K104 via PsbV-I45/K134 as branch B (Fig. 5A, B). Here we calculated channel radii of branches A and B in the PsbO/V/U-reconstituted and native PSII structures. The radius of branch A was broadened around PsbV-K47 in the reconstituted PSII compared with the native PSII (in the area that is enclosed by dashed vertical lines in Fig. 5C), adopting a partially meandering path toward PsbV-K47. In contrast, there was almost no change in the radius of branch B between the reconstituted and native PSII structures (Fig. 5D). These results suggest that branch A within the O1 channel may contribute to water delivery during the S_2_ → S_3_ and S_3_ → S_0_ transitions.

**Figure 5.**
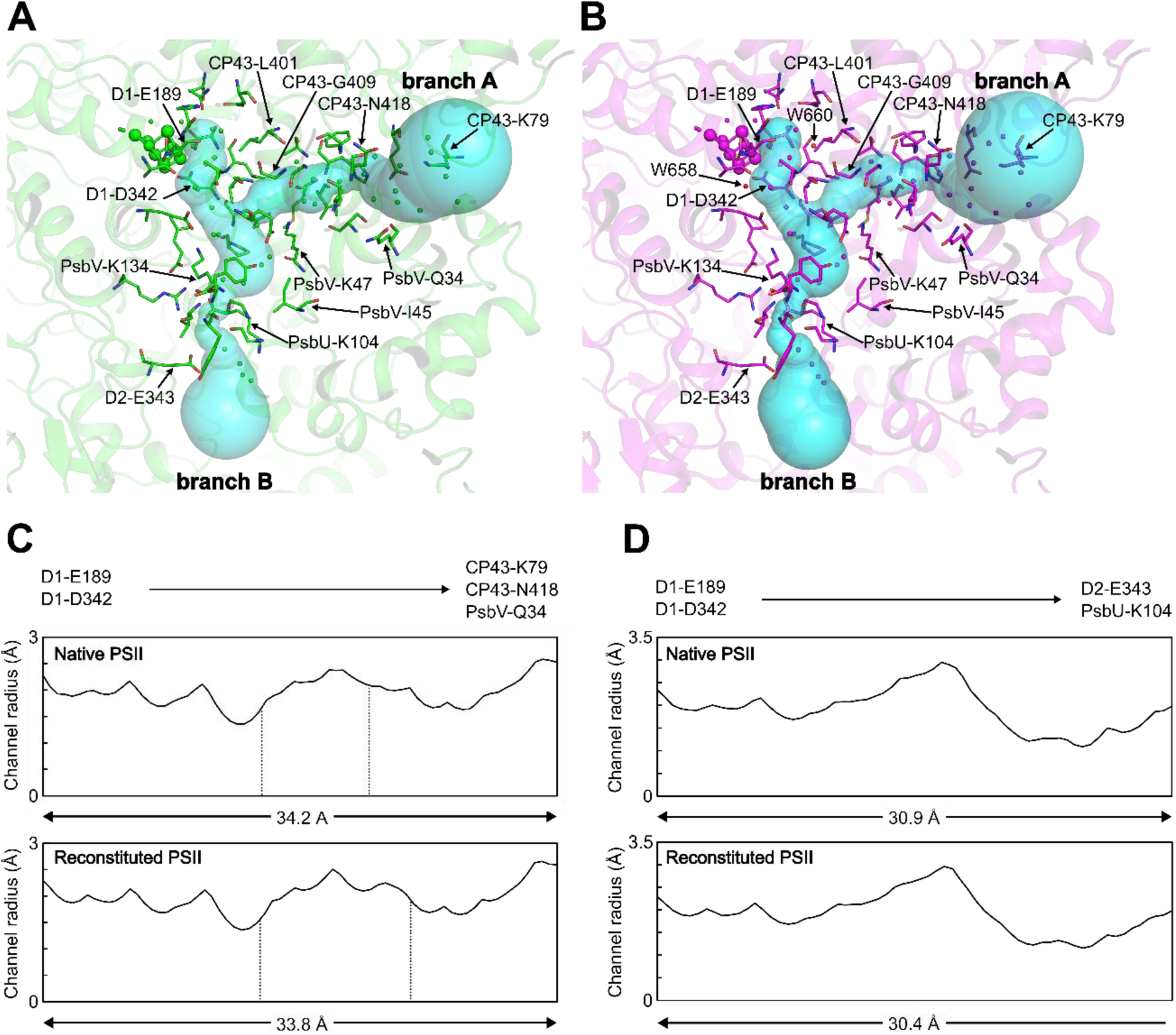
Water networks in the O1 channel in the A-monomers. (**A, B**) Structures of proteins and cofactors in the branches A and B of the O1 channel in the A-monomer. The PsbO/V/U-reconstituted and native PSII structures are colored green (**A**) and magenta (**B**), respectively. The cyan surface models show the cavity of the branches A and B calculated by the CAVER program. (**C, D**) Distribution of the channel radii of the branches A (**C**) and B (**D**) in the native (upper panels) and reconstituted (lower panels) PSII. Branch A is from D1-E189/D342 to CP43-K79/N418 and PsbV-Q34 (**C**), and branch B is from D1-E189/D342 to D2-E343 and PsbU-K104 (**D**). The areas surrounded by two vertical, dashed lines indicate the channel region where PsbV-K47 is involved.

A similar disappearance of the corresponding water molecules (W636 and W639, equivalent to W660 and W658, respectively) was also observed in the B-monomer, accompanied by analogous rearrangements of the hydrogen-bond network and reorientation of D1-N298 (Fig. S6). Channel analysis further showed similar features in the B-monomer to those observed in the A-monomer (Fig. S7A–D). In contrast, the native PSII exhibited distinct O1-channel architectures between the A- and B-monomers, with noticeable differences in channel radii (Fig. S7B, C). These observations indicate that reconstitution induces a structural symmetrization of the O1 channel, leading to equivalent hydration states in both monomers, which may in turn give rise to the decreased oxygen-evolving activity in the reconstituted PSII.

### No changes in the Cl-1 channel

The Cl-1 channel is separated into two branches toward the lumenal side: one is from D1-N181/S169 to D1-P66 and D2-E310 via D1-I60/D61 as branch A, and the other one is from D1-N181/S169 to CP47-T434 and PsbO-A177 via D1-E333 and D2-L321 as branch B (Fig. S8A, B). The channel radii of branches A and B within the Cl-1 channel resembled closely between the reconstituted and native PSII structures (Fig. S8C, D).

It has been suggested that the Cl-1 channel serves as a proton transfer pathway during the S_2_ → S_3_ and S_3_ → S_0_ transitions, as investigated by Cl^−^ depletion studies (*67–71*), mutagenesis studies of D1-D61, D2-K317, and the D1-E65/D1-R334/ D2-E312 triad (*61, 63–65, 72, 73*), and theoretical studies (*55, 60, 62, 65, 74*). Recently, QM/MM calculations showed that the D1-E63/D2-E312 dyad function as a gate of proton transfer in the Cl-1 channel by regulating its closed and open forms (*75*). Thus, while the Cl-1 channel is considered to function as a proton transfer pathway from various lines of evidence, TRIR analysis leaves open regarding the possibility that a proton during the S_2_ → S_3_ transition may be transferred through the O1 channel under some specific conditions (*59, 73*). These experiments showed that proton transfer does not completely cease even when the Cl-1 channel is blocked or its efficiency is reduced upon mutations of amino acid residues in the Cl-1 channel or substitution of Cl^−^. This possibility has been further supported by theoretical calculations (*50, 53, 76, 77*). In this study, we showed almost no changes in the branches A and B between the reconstituted and native PSII (Fig. S8). These observations imply no effects of proton release during the S_2_ → S_3_ and S_3_ → S_0_ transitions in the PsbO/V/U-reconstituted PSII.

### No changes in the O4 channel

The radii of the O4 channel were calculated in the PsbO/V/U-reconstituted and native PSII structures (Fig. S9A, B). The radii from D1-D61 and CP43-E354 to PsbU-Y21/N31 in the O4 channel in the reconstituted PSII were virtually identical to those in the native PSII (Fig. S9C).

The O4 channel is a linear water chain from the Mn_4_CaO_5_ cluster to the lumen. The Ala mutation of D1-S169, which interacts with the first water molecule at the O4 site, showed only minor effects on the efficiencies and the kinetics of the S_2_ → S_3_ and S_3_ → S_0_ transitions (*78*) as well as on the water vibrations (*79*). The O4 channel may serve as a proton transfer pathway during the S_0_ → S_1_ transition (*78, 80–83*), as supported by theoretical studies suggesting that this channel is not suitable for water delivery but rather serves as a proton transfer pathway (*50, 74, 80*). The same channel radii of the O4 channel in the reconstituted and native PSII observed in the present study implies no effect of proton transfer during the S_0_ → S_1_ transition in the PsbO/V/U-reconstituted PSII.

### Molecular mechanisms underlying the reduction of oxygen-evolving activity

This study demonstrates the structural changes in amino acid residues and cofactors in the PsbO/V/U-reconstituted PSII compared with the native PSII. The proper rebinding of PsbO, PsbV, and PsbU to PSII was verified by X-ray crystallography (Fig. 1), yielding an overall RMSD value of 0.172 Å between the reconstituted and native structures. The overall geometry of the Mn_4_CaO_5_ cluster and the inter-cofactor distances are largely conserved between the reconstituted and native PSII (Fig. 2, Fig. S2, S3), although a statistically significant shortening of the Mn2–O2 bond was detected (Table S3), suggesting a subtle local rearrangement within the OEC. Despite the restoration of the extrinsic proteins, the oxygen-evolving activity of the reconstituted PSII remains incompletely recovered, reaching only 80–87% of the native level (*33, 34*). This partial recovery is plausibly associated with structural perturbations at both the donor and acceptor sides of PSII upon release-reconstitution of PsbO, PsbV, and PsbU. Indeed, the present structural analysis revealed such perturbations. First, noticeable alterations were observed in the bicarbonate and its surrounding hydrogen-bond networks (Fig. 3, Fig. S4), which may retard electron transfer from Q_A_ to Q_B_. Furthermore, the loss of W658/W660 and rearrangement of the hydrogen-bond network in the O1 channel (Fig. 4, 5, Fig. S7) may contribute to altered water delivery through the O1 channel during the S_2_ → S_3_ and S_3_ → S_0_ transitions. By contrast, proton transfer during the S_2_ → S_3_, S_3_ → S_0_, and S_0_ → S_1_ transitions appears unaffected (Fig. S8, S9). A schematic representation of these findings and their relation to the S-state cycle is shown in Fig. 6. Collectively, these intertwined reactions—encompassing disrupted water delivery, altered electron transfer, and minor rearrangements within the Mn_4_CaO_5_ cluster—provide a structural framework for addressing the long-standing question of why oxygen-evolving activity decreases in extrinsic-protein-reconstituted PSII, and more broadly, for understanding the mechanisms governing electron transfer, water delivery, and proton egress in the photosynthetic water-oxidation reaction.

**Figure 6.**
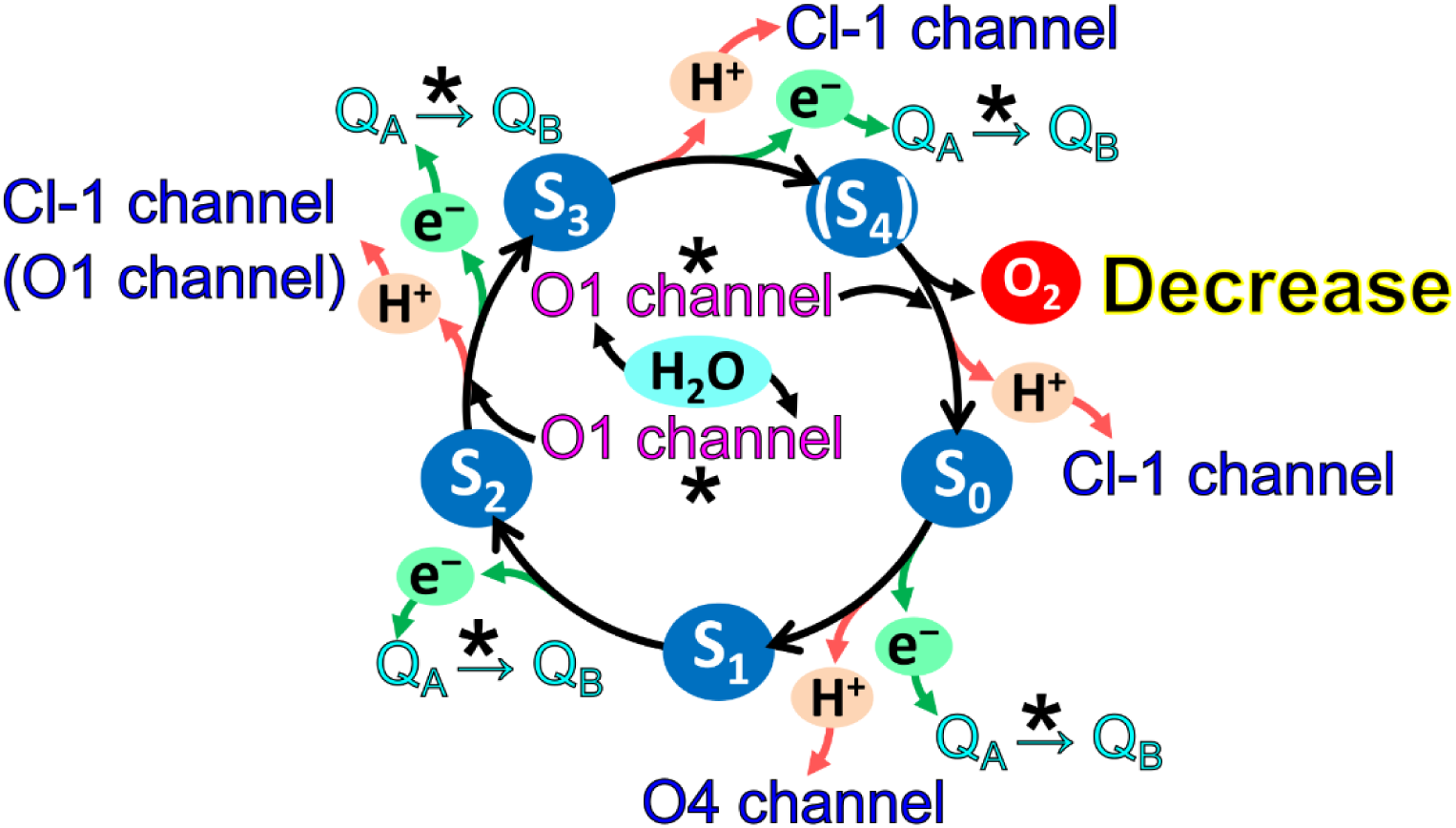
Schematic model for the S-state cycle in the PsbO/V/U-reconstituted PSII. Asterisks indicate the possible retardation processes of water delivery in the O1 channel and electron transfer from Q_A_ to Q_B_ in the reconstituted PSII.

## Methods

### Purification of PSII cores and extrinsic subunits, and reconstitution experiments

The thermophilic cyanobacterium *T. vulcanus* NIES-2134 was grown and its PSII dimer cores were purified as described previously (*31, 32*). The extrinsic PsbO, PsbV, and PsbU subunits were removed from PSII with the treatment of 1 M CaCl_2_ as described previously (*33, 84*), and subsequently dialyzed with 20 mM Mes-NaOH (pH 6.5) at 4 °C. The PSII cores depleted of PsbO, PsbV, and PsbU were reconstituted with all the extrinsic proteins at a molar ratio of 1:3 (PSII:extrinsic proteins), according to the previous methods (*33, 84*). The protein compositions of the PSII samples were analyzed by SDS-PAGE following the method of Ikeuchi and Inoue (*85*).

### Crystallization and structural analysis

The PsbO/V/U-reconstituted PSII complexes were crystallized as described previously (*31, 32, 86*). Briefly, crystals of the PsbO/V/U-reconstituted PSIIs grew to a size of around 0.8 × 0.4 × 0.2 mm, and they were collected and treated by cryoprotectant solution. X-ray diffraction images were collected at the beamline BL41XU of SPring-8, Japan. The dataset obtained was indexed, integrated, and scaled with XDS (*87*). The initial phase up to a resolution of 4.0 Å was obtained by molecular replacement with Phaser MR in CCP4 program suite (*88*) using the 1.9-Å resolution structure of native PSII (PDB accession code: 3WU2) as the search model. The structure was refined to a resolution of 2.0 Å with REFMAC5 in the CCP4 suite (*89*) and Phenix refinement (*90*). The RMSD was calculated using LSQKAB in the CCP4 suite (*91*). Model building was performed with Coot (*92*), and figures were made with Chimera (*93*) and PyMOL (*94*).

Channel calculations were performed using the CAVER v3.03 PyMOL Plugin (*95*), with cavities formed by amino acid residues near the Mn_4_O_5_Ca cluster selected as starting point for the analysis of previously reported water channels (*96*). The CAVER program was executed with the probe radius, shell radius, and shell depth configured to 0.7, 3.0, and 4.0 Å, respectively. Water molecules were excluded for channel calculations. A detailed analysis of the channel radii and the contributions of specific amino acids to channel formation was performed using CAVER Analyst 2.0 (*97*).

### Estimation of coordinate errors for interatomic distances around the bicarbonate molecule and within the OEC

The uncertainties of interatomic distances around the bicarbonate molecule and within the OEC were evaluated based on the diffraction-component precision index (DPI) (*98, 99*) and the RMSD. The distance error between two structures derived from DPI, denoted as σ_DPI_, was defined as:

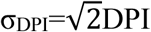

The factor of the square root of two accounts for the propagation of independent coordinate uncertainties from the two structures being compared. Because DPI assumes that all atoms in the structure share an identical temperature factor, it tends to overestimate coordinate errors in regions with low average B-factors, such as the OEC. To correct for this, we additionally estimated the coordinate uncertainty around the OEC using a previously described method (*100*). Specifically, the RMSD between the two structures was calculated using regions with the lowest average B-factors, comprising Cα atoms of D1-F17–S222 and S268–A344, and D2-G34–T221 and T243–L352. The distance error derived from RMSD, denoted as σ_RMSD_, was defined as:

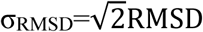

The RMSD-based and DPI-based errors were defined as the minimum and maximum estimates, respectively, of the coordinate uncertainty around the OEC. The midpoint between these two values (0.132 Å) was adopted as the representative coordinate error for the OEC region (Table S2) and used to assess the reliability and significance of changes in interatomic distances between the two structures. Statistical confidence was evaluated using a two-tailed test based on the normal distribution, and differences with a confidence level greater than 95% (approximately 2σ) were considered statistically significant.

## Data availability

The structure reported in this article has been deposited in PDB with the accession code of 9LP8. The restraint and geometry files used for the refinement of the OEC structure are available from the Zenodo data repository (https:// doi.org/10.5281/zenodo.17854142). All other data are available from the corresponding authors upon reasonable request.

## Acknowledgments

We thank the staff members at the beamline BL41XU of SPring-8 for their assistance in data collection under the proposal number 2021B2741. This work was supported by JSPS KAKENHI grant Nos. JP23K14211 (Y.N.), JP22H04916 (J.-R.S.), and JP23H02423 (R.N.); and JCC-Saneyoshi Scholarship Foundation (R.N.).

## Authorship contributions

R.N. conceived the project; Y.N. and R.N. prepared PSII samples; Y.N. performed X-ray crystal structure analysis; Y.N., K.K., and R.N. analyzed structure; Y.N. and R.N. drafted the original manuscript; J.-R.S. modified the manuscript; R.N. wrote the final manuscript; and all of the authors contributed to the interpretations of the results and improvement of the manuscript.

