## Supplementary information for "Structural basis for impaired oxygen evolution in extrinsic-protein-reconstituted photosystem II"

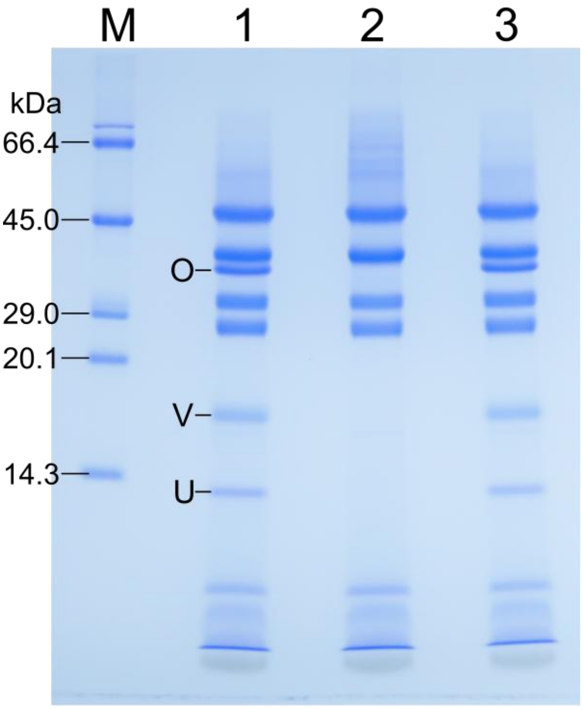


**Figure S1 | Evidence for the release and reconstitution of the three extrinsic proteins.**
The presence or absence of PsbO, PsbV, and PsbU was assessed by SDS-PAGE. Lane M, molecular weight marker; lane 1, untreated PSII; lane 2, PSII depleted of the three extrinsic proteins; lane 3, PSII reconstituted with the three extrinsic proteins. PsbO, PsbV, and PsbU are labeled as O, V, and U, respectively. The corresponding protein bands were detected in lanes 1 and 3.

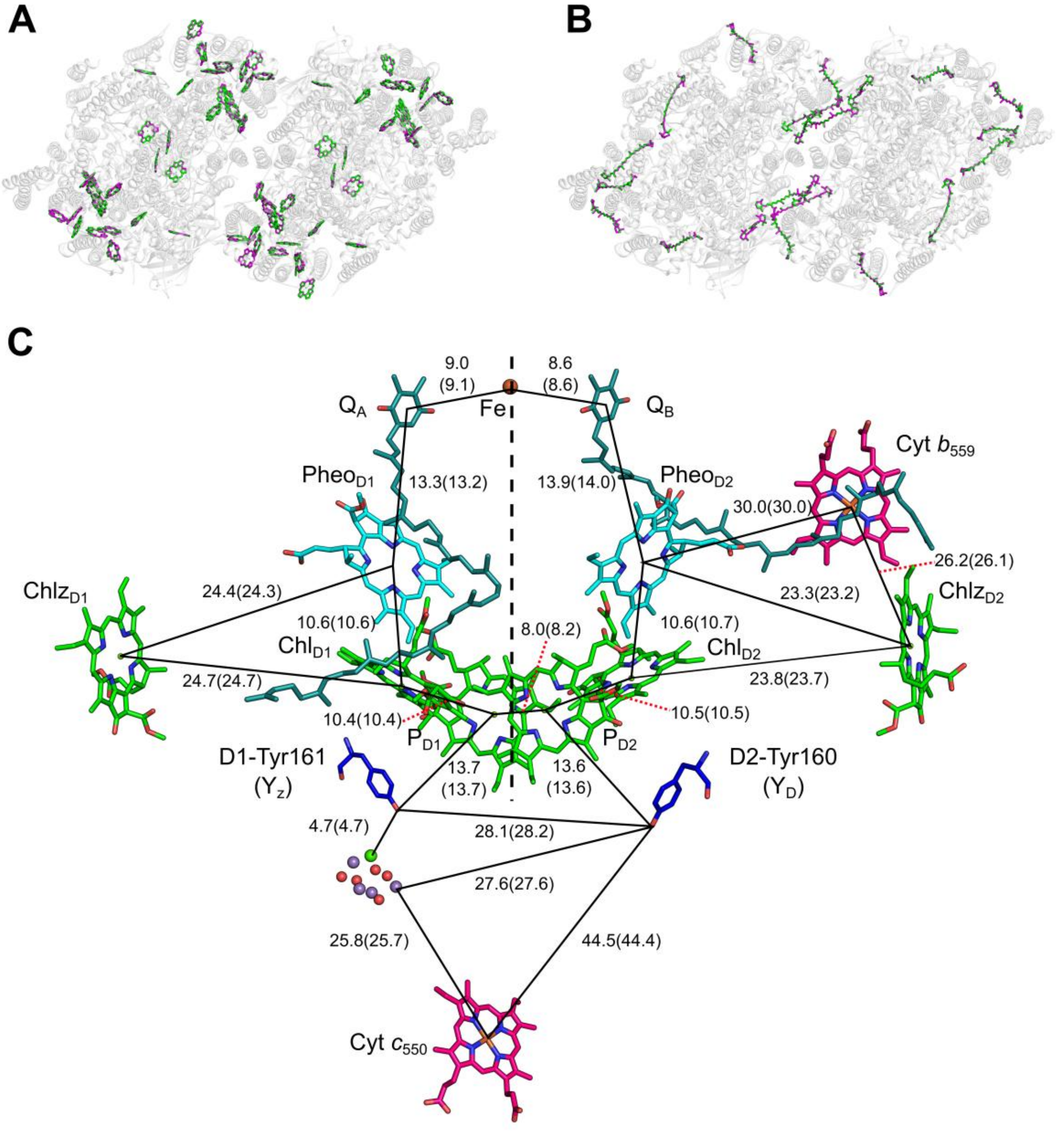


**Figure S2 | Arrangements of pigments and cofactors in the reconstituted and native PSII cores.**

(**A, B**) Structures viewed from the cytosolic side. Chls (**A**) and Cars (**B**) are colored, and proteins are shown in grey. Only rings of the Chl molecules are depicted. The PsbO/V/U-reconstituted and native PSII structures are colored green and magenta, respectively. (**C**) Comparisons among redox-active cofactors in the A-monomers of the native and reconstituted PSII. Interactions are indicated by lines, and the numbers are distances in Å. The distances of the reconstituted PSII are shown as numbers, and those of the native PSII are shown as numbers in parentheses.

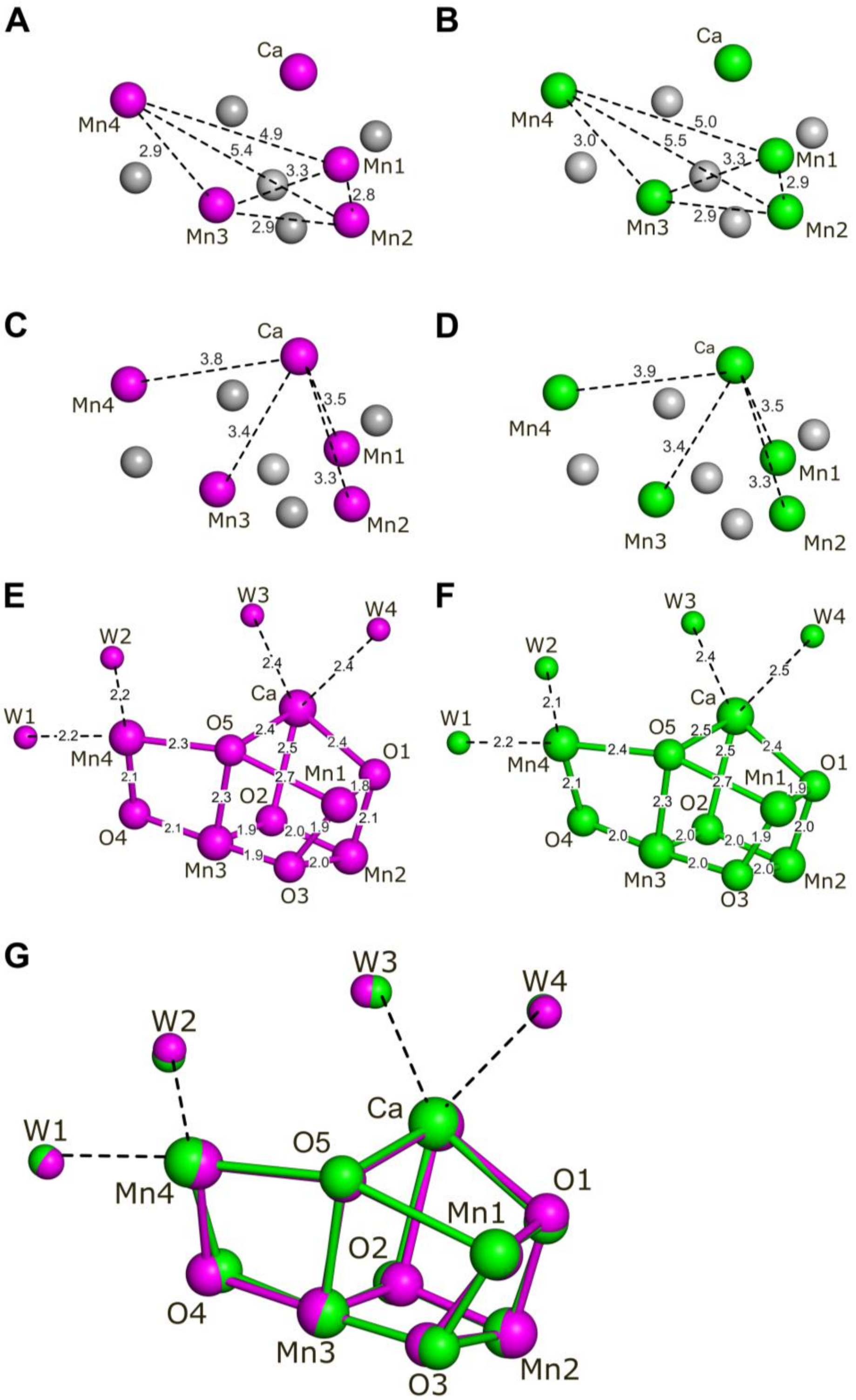


**Figure S3 | Structural comparison of the OEC in the B-monomers of the native and reconstituted PSII.**

(**A, B**) Distances between Mn atoms. (**C, D**) Distances between Mn and Ca atoms. (**E, F**) Distances between the $Mn_4CaO_5$ cluster and its coordinating water molecules. (**G**) Superposition of the OECs from the two PSII structures. All panels show the OECs in the B-monomers of the native PSII (magenta; PDB 3WU2) and the PsbO/V/U-reconstituted PSII (green).

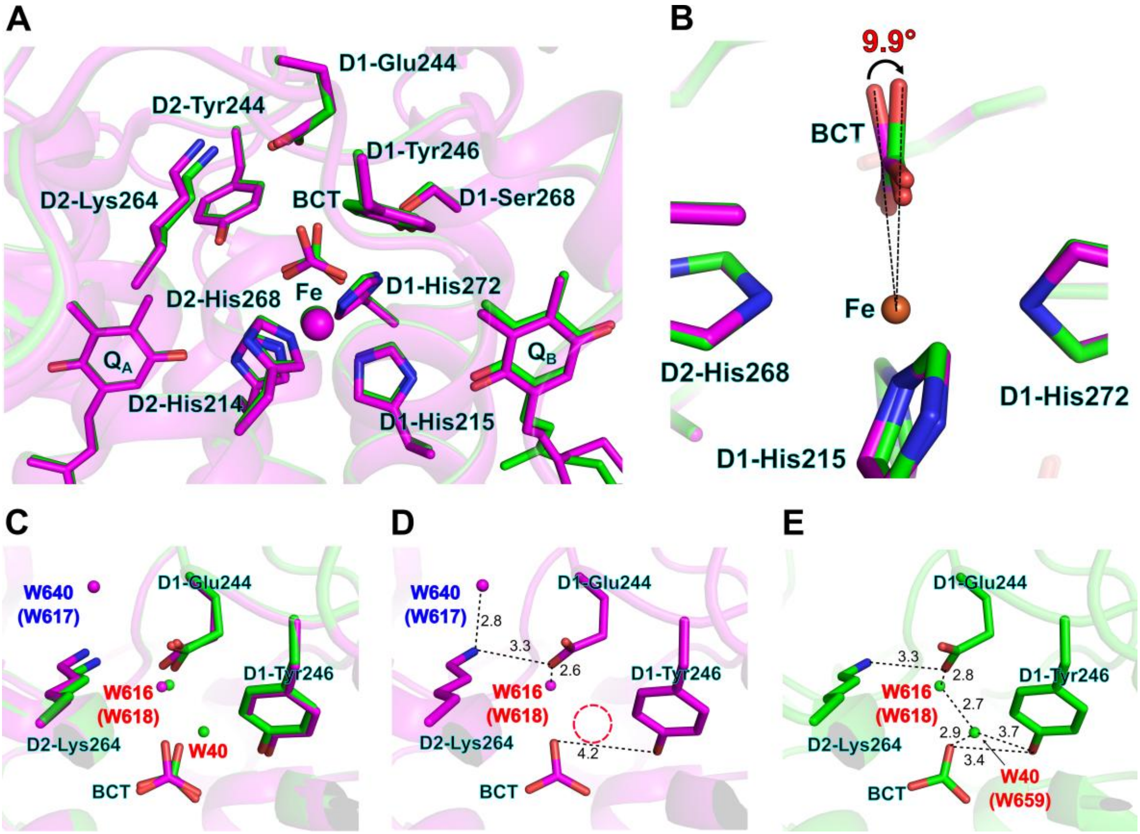


**Figure S4 | Structural comparison of the acceptor side in the B-monomers of the native and reconstituted PSII.**

(**A**) Structures around the bicarbonate (BCT) molecule and non-heme iron (Fe). The PsbO/V/U-reconstituted and native PSII structures are colored green and magenta, respectively. (**B**) Angle of BCT. The angle was calculated from three points: O1 of BCT in the native PSII (PDB 3WU2), Fe, and O1 of BCT in the reconstituted PSII, based on superposition of the Fe atom between the two structures. (**C–E**) Superposition of BCT and its nearby amino acid residues and water molecules between the structures of reconstituted and native PSII (**C**), structure of the reconstituted PSII (**D**), and structure of the native PSII (**E**). Interactions are indicated by dashed lines, and the numbers are distances in Å. Water molecules of the D1 and D2 subunits are labeled with red and blue colors, respectively; numbers in parentheses indicate the corresponding water molecule in the A-monomer. A newly observed D1-derived water molecule, designated as W40 (chain W) in the deposited structure, was not present in the native PSII.

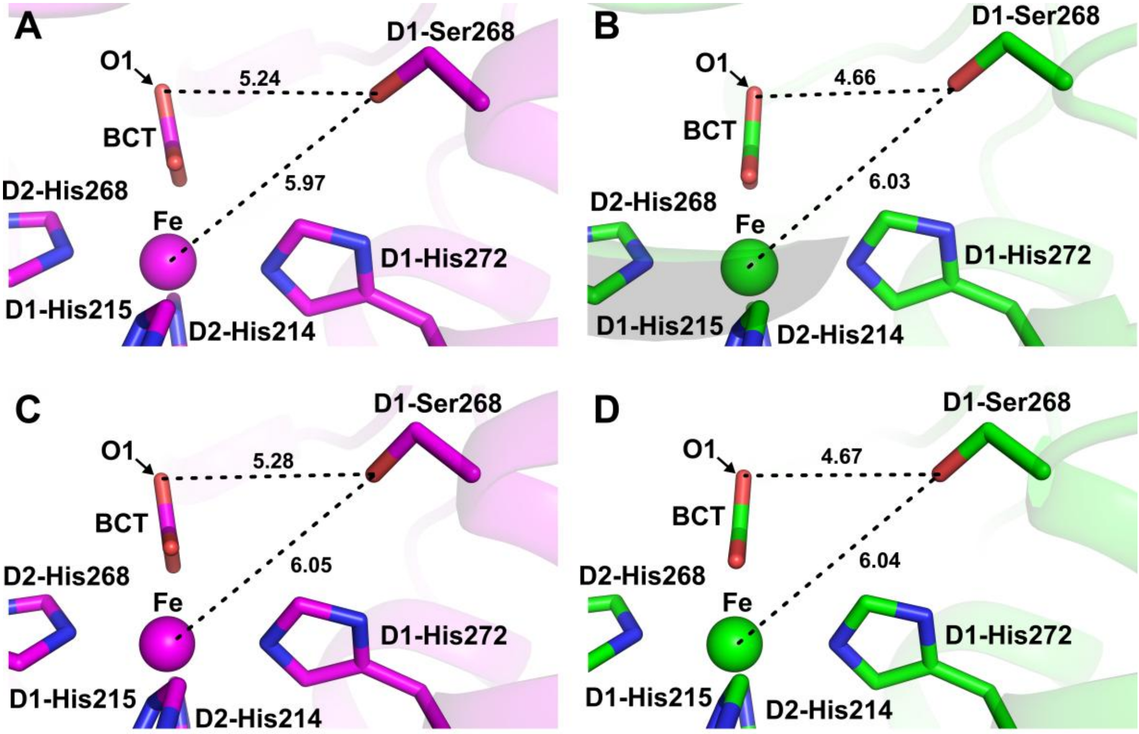


**Figure S5 | Structural comparison of the bicarbonate-binding site in the native and reconstituted PSII.**

Structures around the bicarbonate (BCT) molecule, non-heme iron (Fe), and amino acid residues in the A-monomers (**A, C**) and B-monomers (**B, D**) in the native PSII (magenta; PDB 3WU2) and the PsbO/V/U-reconstituted PSII (green). Interactions are indicated by dashed lines, and the numbers are distances in Å.

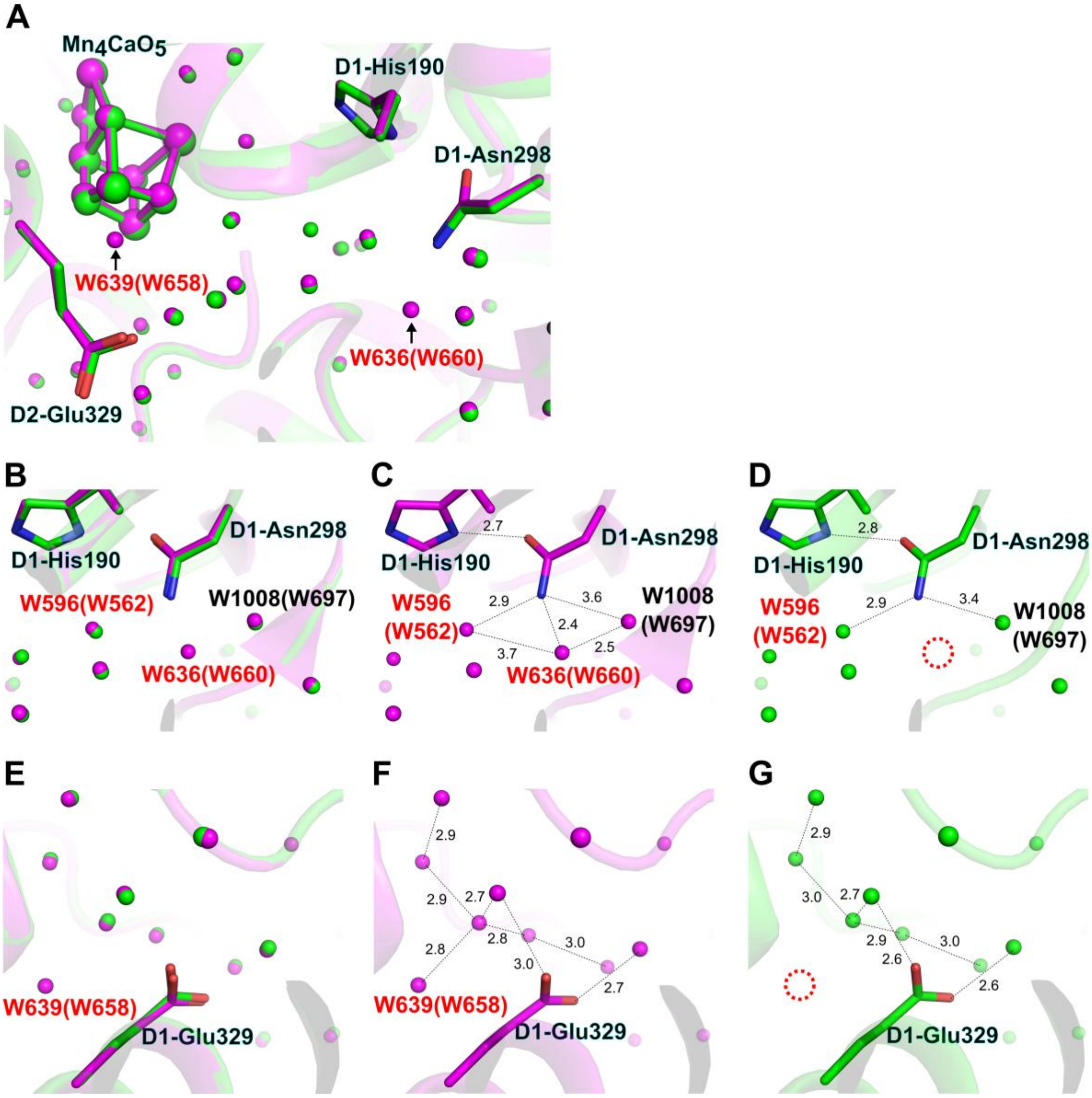


**Figure S6 | Structural comparison of the O1 channel in the B-monomers of the native and reconstituted PSII.**

(**A**) Structures of proteins and cofactors in the O1 channel. The PsbO/V/U-reconstituted and native PSII structures are colored green and magenta, respectively. (**B–D**) Interactions of W636 with proteins and water molecules in the structures of the reconstituted and native PSII superposed (**B**), the native PSII (**C**), and the reconstituted PSII (**D**). (**E–G**) Interactions of W639 with protein and water molecules in the structures of the reconstituted and native PSII superposed (**E**), the native PSII (**F**), and the reconstituted PSII (**G**). Interactions are indicated by dashed lines, and the numbers are distances in Å. Water molecules of the D1 and CP43 subunits are labeled with red and black colors, respectively. Areas enclosed by red dashed circles indicate the absence of W636 (**D**) and W639 (**G**) in the reconstituted PSII. Numbers in parentheses indicate the corresponding water molecule in the A-monomer.

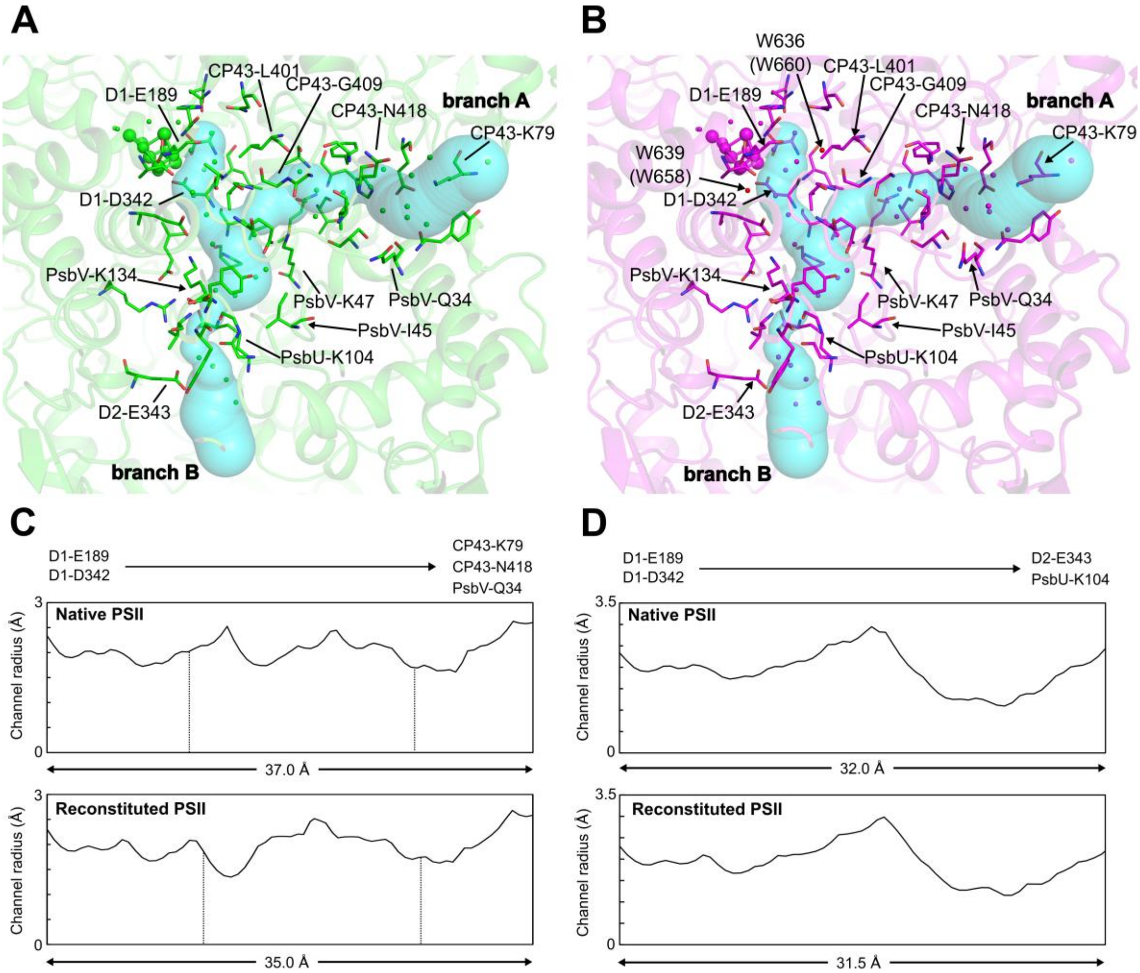


**Figure S7 | Water networks in the O1 channel in the B-monomers.**

(**A, B**) Structures of proteins and cofactors in the branches A and B of the O1 channel in the B-monomer. The PsbO/V/U-reconstituted and native PSII structures are colored green (**A**) and magenta (**B**), respectively. The cyan surface models show the cavity of the branches A and B calculated by the CAVER program. (**C, D**) Distribution of the channel radii of the branches A (**C**) and B (**D**) in the native (upper panels) and reconstituted (lower panels) PSII. Branch A is from D1-E189/D342 to CP43-K79/N418 and PsbV-Q34 (**C**), and branch B is from D1-E189/D342 to D2-E343 and PsbU-K104 (**D**). The areas surrounded by two vertical, dashed lines indicate the channel region where PsbV-K47 is involved. Numbers in parentheses indicate the corresponding water molecule in the A-monomer.

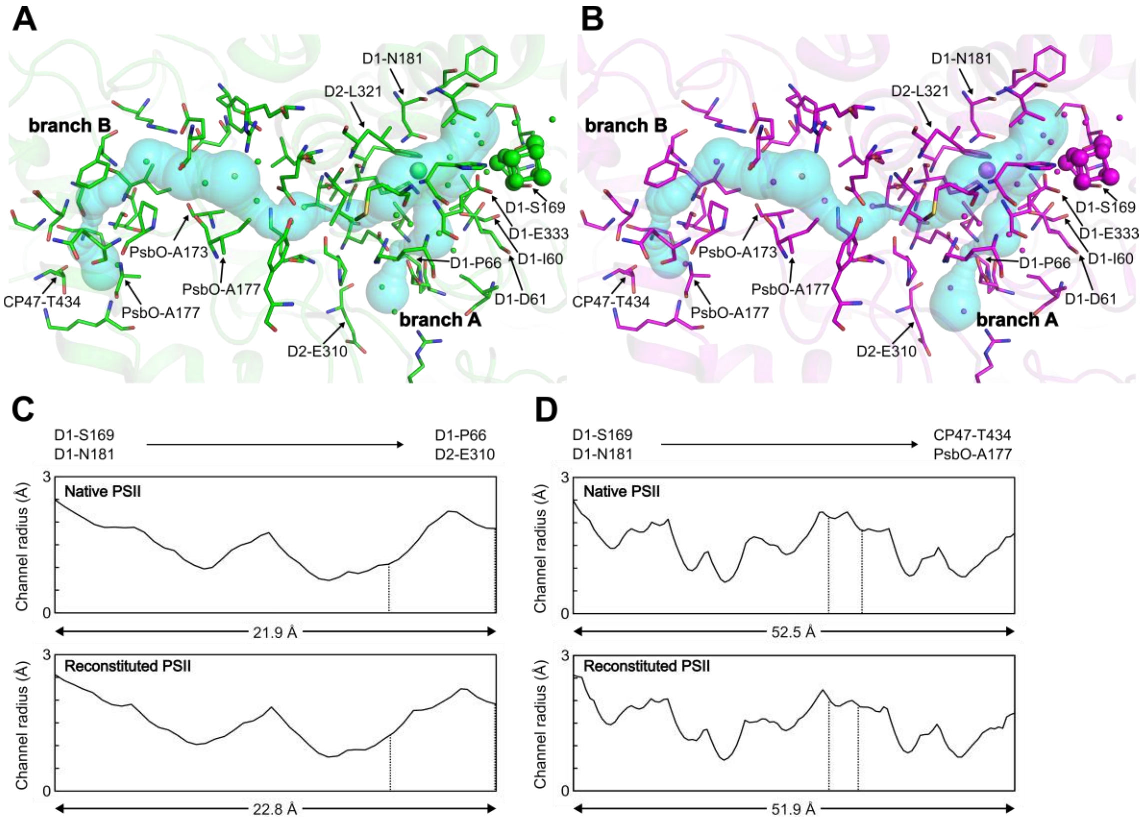


**Figure S8 | Water network in the Cl-1 channel.**

(**A, B**) Structures of proteins and cofactors in branches A and B of the Cl-1 channel of the A-monomers. The PsbO/V/U-reconstituted and native PSII structures are colored green (**A**) and magenta (**B**), respectively. The cyan surface models show the cavity of branches A and B calculated by the CAVER program. (**C, D**) Distribution of the channel radii of branches A (**C**) and B (**D**) in the native (upper panels) and reconstituted (lower panels) PSII. Branch A is from D1-N181/S169 to D1-P66 and D2-E310 (**C**), and branch B is from D1-N181/S169 to CP47-T434 and PsbO-A177 (**D**). The areas surrounded by vertical, dashed lines in panels **C** and **D** indicate the channel regions where D2-E310 and PsbO-I172/A173 are involved, respectively.

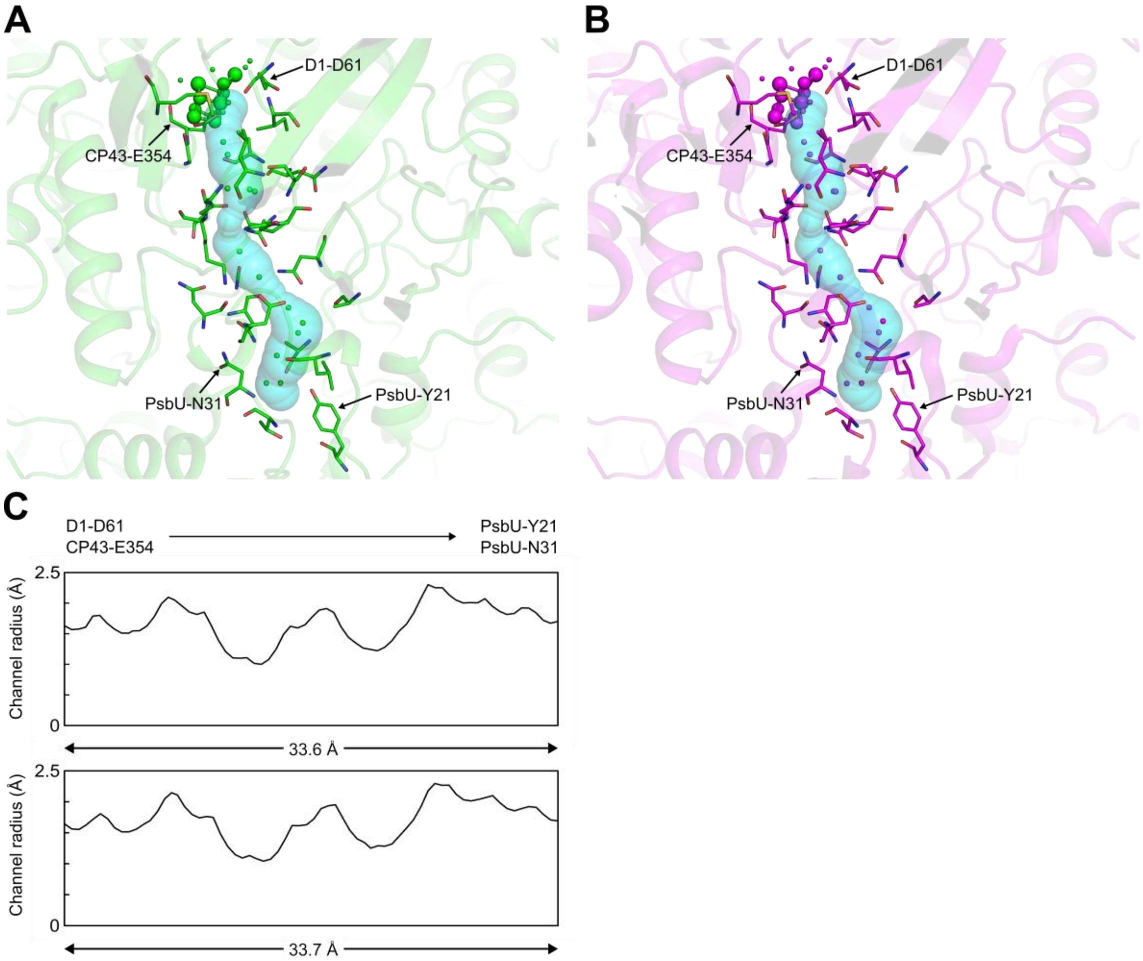


**Figure S9 | Water network in the O4 channel.**

(**A, B**) Structures of proteins and cofactors in the O4 channel of the A-monomers. The PsbO/V/U-reconstituted and native PSII structures are colored green (**A**) and magenta (**B**), respectively. The cyan surface models show the cavity of the O4 channel calculated by the CAVER program. (**C**) Distribution of the channel radii of the O4 channel in the native (upper panel) and reconstituted (lower panel) PSII. The O4 channel is from D1-D61 and CP43-E354 to PsbU-T21/N31.

**Table S1 | Data collection and refinement statistics.**

| | PsbO/V/U-reconstituted PSII |
|---|---|
| **Data collection statistics** | |
| Wavelength (Å) | 1.00 |
| Resolution range | 49.1 – 2.00 (2.07 – 2.00)[a] |
| Space group | $P2_12_12_1$ |
| Unit cell (Å) | a=123.3, b=228.9, c=286.8 |
| Total reflections | 4,161,498 (418,440)[a] |
| Unique reflections | 542,391 (53,767)[a] |
| Multiplicity | 7.7 (7.8)[a] |
| Completeness (%) | 99.95 (99.96)[a] |
| Mean I/σ (I) | 23.52 (2.18)[a] |
| $R_{merge}$ | 0.050 (0.906)[a] |
| $R_{pim}$ | 0.019 (0.344)[a] |
| $CC_{1/2}$ | 0.999 (0.848)[a] |
| **Refinement statistics** | |
| Resolution (Å) | 49.1–2.00 |
| $R_{work}/R_{free}$ (%) | 0.152 /0.178 |
| No. of protein residues | 5,273 |
| No. of water molecules | 2,750 |
| Wilson B-factor (Å$^2$) | 39.1 |
| R. m. s. d. bond length (Å) | 0.02 |
| R. m. s. d. bond angles (°) | 1.80 |
| Ramachandran plot (%)[b] | |
| Favored | 98.2 |
| Allowed | 1.7 |
| Outliers | 0.1 |
| Rotamer outliers (%) | 0.69 |
| Clashscore | 4.37 |
| Average B-factor (Å$^2$) | |
| overall | 51.0 |
| protein | 49.4 |
| ligands | 56.5 |

[a]Statistics for the highest-resolution shell are shown in parentheses.
[b]Ramachandran plot was calculated by MolProbity.

**Table S2 | Calculated errors in atomic coordinates and interatomic distances.**

| | Native PSII (3WU2) | PsbO/V/U-reconstituted PSII |
|---|---|---|
| RMSD | 0.075 | |
| $\sigma_{RMSD}$ | 0.106 | |
| DPI | 0.111 | 0.113 |
| $\sigma_{DPI}$ | 0.158 | 0.159 |
| Interatomic distance error of OEC | 0.132 | |

**Table S3 | Interatomic distances of the OEC.**

| | | | Native (3WU2) | | | | PsbO/V/U-reconstituted PSII | | | | A–A / B–B monomer comparison | | |
|---|---|---|---|---|---|---|---|---|---|---|---|---|---|
| | | | A | B | average | A-B | A | B | average | A-B | A-A | B-B | average-diff |
| Mn-Mn | | | | | | | | | | | | | |
| Mn1 | - | Mn2 | 2.87 | 2.76 | 2.82 | 0.11 | 2.87 | 2.86 | 2.87 | 0.01 | 0.00 | 0.10 | 0.05 |
| Mn1 | - | Mn3 | 3.28 | 3.29 | 3.29 | 0.01 | 3.28 | 3.29 | 3.29 | 0.01 | 0.00 | 0.00 | 0.00 |
| Mn1 | - | Mn4 | 4.99 | 4.92 | 4.96 | 0.07 | 5.02 | 5.03 | 5.03 | 0.01 | 0.03 | 0.11 | 0.07 |
| Mn2 | - | Mn3 | 2.88 | 2.92 | 2.90 | 0.04 | 2.87 | 2.85 | 2.86 | 0.02 | 0.01 | 0.07 | 0.04 |
| Mn2 | - | Mn4 | 5.44 | 5.38 | 5.41 | 0.06 | 5.47 | 5.47 | 5.47 | 0.00 | 0.03 | 0.09 | 0.06 |
| Mn3 | - | Mn4 | 2.97 | 2.89 | 2.93 | 0.08 | 3.02 | 3.02 | 3.02 | 0.00 | 0.05 | 0.13 | 0.09 |
| Mn-Ca | | | | | | | | | | | | | |
| Ca | - | Mn1 | 3.55 | 3.49 | 3.52 | 0.06 | 3.60 | 3.51 | 3.56 | 0.09 | 0.05 | 0.02 | 0.04 |
| Ca | - | Mn2 | 3.39 | 3.30 | 3.35 | 0.09 | 3.41 | 3.31 | 3.36 | 0.10 | 0.02 | 0.01 | 0.01 |
| Ca | - | Mn3 | 3.42 | 3.41 | 3.42 | 0.01 | 3.47 | 3.39 | 3.43 | 0.08 | 0.05 | 0.02 | 0.01 |
| Ca | - | Mn4 | 3.79 | 3.80 | 3.80 | 0.01 | 3.85 | 3.87 | 3.86 | 0.02 | 0.06 | 0.07 | 0.06 |
| Mn-O | | | | | | | | | | | | | |
| Mn1 | - | O1 | 1.85 | 1.80 | 1.83 | 0.05 | 2.00 | 1.91 | 1.96 | 0.09 | 0.15 | 0.11 | 0.13 |
| Mn1 | - | O3 | 1.84 | 1.93 | 1.89 | 0.09 | 1.87 | 1.86 | 1.87 | 0.01 | 0.03 | 0.07 | 0.02 |
| Mn1 | - | O5 | 2.65 | 2.66 | 2.66 | 0.01 | 2.66 | 2.67 | 2.67 | 0.01 | 0.01 | 0.01 | 0.01 |
| Mn2 | - | O1 | 2.08 | 2.07 | 2.08 | 0.01 | 2.02 | 1.97 | 2.00 | 0.05 | 0.06 | 0.10 | 0.08 |
| Mn2 | - | O2 | 2.16 | 1.96 | 2.08 | 0.20 | 1.90 | 2.03 | 1.97 | 0.13 | 0.26 | 0.07 | 0.11 |
| Mn2 | - | O3 | 2.13 | 2.02 | 2.08 | 0.11 | 2.01 | 2.03 | 2.02 | 0.02 | 0.12 | 0.01 | 0.06 |
| Mn3 | - | O2 | 1.93 | 1.91 | 1.92 | 0.02 | 1.96 | 1.95 | 1.96 | 0.01 | 0.03 | 0.04 | 0.04 |
| Mn3 | - | O3 | 2.24 | 1.86 | 2.05 | 0.38 | 2.12 | 2.04 | 2.08 | 0.08 | 0.12 | 0.18 | 0.03 |
| Mn3 | - | O4 | 2.05 | 2.08 | 2.08 | 0.03 | 1.96 | 1.98 | 1.97 | 0.02 | 0.09 | 0.10 | 0.11 |
| Mn4 | - | O3 | 2.28 | 2.25 | 2.27 | 0.03 | 2.22 | 2.31 | 2.27 | 0.09 | 0.06 | 0.06 | 0.00 |
| Mn4 | - | O4 | 2.09 | 2.05 | 2.07 | 0.04 | 2.05 | 2.13 | 2.09 | 0.08 | 0.04 | 0.08 | 0.02 |
| Mn4 | - | O5 | 2.43 | 2.34 | 2.39 | 0.09 | 2.42 | 2.43 | 2.43 | 0.01 | 0.01 | 0.09 | 0.04 |
| Ca-O | | | | | | | | | | | | | |
| Ca | - | O1 | 2.39 | 2.44 | 2.42 | 0.05 | 2.41 | 2.46 | 2.44 | 0.05 | 0.02 | 0.02 | 0.02 |
| Ca | - | O2 | 2.53 | 2.48 | 2.51 | 0.05 | 2.50 | 2.54 | 2.52 | 0.04 | 0.03 | 0.06 | 0.01 |
| Ca | - | O5 | 2.43 | 2.57 | 2.50 | 0.14 | 2.51 | 2.56 | 2.54 | 0.05 | 0.08 | 0.01 | 0.04 |
| Water | | | | | | | | | | | | | |
| Mn4 | - | W1 | 2.27 | 2.17 | 2.22 | 0.10 | 2.24 | 2.18 | 2.21 | 0.06 | 0.03 | 0.01 | 0.01 |
| Mn4 | - | W2 | 2.12 | 2.20 | 2.16 | 0.08 | 2.10 | 2.10 | 2.10 | 0.00 | 0.02 | 0.10 | 0.06 |
| Ca | - | W3 | 2.34 | 2.42 | 2.38 | 0.08 | 2.44 | 2.41 | 2.43 | 0.03 | 0.10 | 0.01 | 0.05 |
| Ca | - | W4 | 2.46 | 2.40 | 2.43 | 0.06 | 2.40 | 2.38 | 2.39 | 0.02 | 0.06 | 0.02 | 0.04 |
| Ligand | | | | | | | | | | | | | |
| Mn1 | - | Glu189 | 1.87 | 1.71 | 1.79 | 0.16 | 1.79 | 1.81 | 1.80 | 0.02 | 0.08 | 0.10 | 0.01 |
| Mn1 | - | His332 | 2.09 | 2.15 | 2.12 | 0.06 | 2.14 | 2.13 | 2.14 | 0.01 | 0.05 | 0.02 | 0.02 |
| Mn1 | - | Asp342 | 2.22 | 2.28 | 2.25 | 0.06 | 2.12 | 2.15 | 2.14 | 0.03 | 0.10 | 0.13 | 0.11 |
| Mn2 | - | Asp342 | 2.15 | 2.04 | 2.10 | 0.11 | 2.11 | 2.15 | 2.13 | 0.04 | 0.04 | 0.11 | 0.03 |
| Mn2 | - | Ala344 | 1.97 | 1.85 | 1.91 | 0.12 | 1.94 | 1.93 | 1.94 | 0.01 | 0.03 | 0.08 | 0.03 |
| Mn2 | - | Glu354 | 2.09 | 2.16 | 2.13 | 0.07 | 2.05 | 2.05 | 2.05 | 0.00 | 0.04 | 0.11 | 0.08 |
| Mn3 | - | Glu333 | 2.02 | 1.94 | 1.98 | 0.08 | 1.97 | 2.00 | 1.99 | 0.03 | 0.05 | 0.06 | 0.01 |
| Mn3 | - | Glu354 | 2.21 | 2.17 | 2.19 | 0.04 | 2.14 | 2.10 | 2.12 | 0.04 | 0.07 | 0.07 | 0.07 |
| Mn4 | - | Asp170 | 2.06 | 2.11 | 2.09 | 0.05 | 2.12 | 2.07 | 2.10 | 0.05 | 0.06 | 0.04 | 0.01 |
| Mn4 | - | Glu333 | 2.17 | 2.13 | 2.15 | 0.04 | 2.11 | 2.08 | 2.10 | 0.03 | 0.06 | 0.05 | 0.05 |
| Ca | - | Asp170 | 2.42 | 2.36 | 2.39 | 0.06 | 2.41 | 2.41 | 2.41 | 0.00 | 0.01 | 0.05 | 0.02 |
| Ca | - | Ala344 | 2.55 | 2.43 | 2.49 | 0.12 | 2.44 | 2.40 | 2.42 | 0.04 | 0.11 | 0.03 | 0.07 |
